# Two-Dimensional Phase Separation of DNA Nanomotifs Anchored to Lipid Bilayers

**DOI:** 10.64898/2026.05.19.724116

**Authors:** Vinavadini Ramnarain, Ana Vazquez, Santiago Labale, Aurélie Di Cicco, Koyomi Nakazawa

## Abstract

Spatial organization and temporal regulation of membrane components are essential for achieving complex functions in artificial cells, such as cell division and signalling. DNA-based molecular tools provide a powerful means to control biomolecular interactions with high precision. Here, we investigate the phase behavior of cholesterol-modified, star-shaped DNA nanomotifs anchored to the lipid bilayers of giant unilamellar vesicles (GUVs), by using fluorescence confocal microscopy and cryo-electron microscopy. These motifs spontaneously anchor to the lipid bilayers via hydrophobic interactions and exhibit distinct spatial organization depending on their sticky end sequences. Motifs with complementary sticky end sequences interact and distribute uniformly, while orthogonal motifs with different sticky end sequences segregate into isolated gel-like domains with limited lateral mobility. Notably, the phase separation of motifs does not require lipid phase separation, indicating that DNA-driven organization can take place independently of lipid phase separation. The behavior of this system is governed by the interplay of three key parameters: (i) hydrophobic anchoring via cholesterol, (ii) electrostatic repulsion between negatively charged DNA nanomotifs, and (iii) sticky end interactions. The observed two-dimensional phase separation of orthogonal DNA nanomotifs at the GUV interface presents a novel strategy for controlling lateral membrane organization in GUV systems. This approach would offer flexibility in membrane composition and enables molecular positioning, thereby achieving a high degree of organization on the surface in artificial cell models.

## Introduction

In recent years, the development of artificial cells has emerged as a powerful approach for studying cellular mechanisms, the origin of life, and future biomedical applications^[1]^. The attempts to recreate cellular functions in cell-mimicking compartments, such as giant unilamellar vesicles (GUVs) and water-in-oil droplets^[2]^, have been made, including the synthesis of ATP and proteins, cytoskeleton growth and metabolic reactions. Achieving complex cellular functions such as division and signalling demands both spatial organization and temporal regulation. However, strategies enabling precise spatiotemporal control of molecular interactions is currently limited in synthetic cell models.

To address spatiotemporal control in artificial cells, the formation of artificial organelles by liquid-liquid phase separation (LLPS) of polymers^[3]^ or microgels^[4]^, or light-based control system^[5]^, have been developed and used to compartmentalize individual biochemical reactions in solutions^[6]^. On the other hand, distinct spatial organization of membrane proteins and lipids at the synthetic cell surfaces is essential for the response to external cues and signalling specificity through localized activation of membrane components, as seen in living systems^[7,8]^. The spatiotemporal control of biomimetic membranes *in vitro* has traditionally relied on two main strategies: the development of photocontrol systems^[5,9,10]^ and the exploitation of phase separations in lipid mixtures^[11–14]^. While photocontrol provides high spatiotemporal precision, it often requires specific optogenetic tools and is constrained by spectral overlap in fluorescence imaging. On the other hand, phase separation of lipid mixtures provides spatial organization but is highly dependent on lipid species which can limit compatibility with membrane proteins or functional components.

DNA-based molecular tools provide a powerful means for facile and precise control in biomolecular coordination, and is used as versatile tool to control biomolecular interactions in various fields^[15–18]^. A recent study from Takinoue group^[19]^ demonstrates that star-shaped DNA nanomotifs undergo LLPS in solution, enabling the formation of a condensate, a membraneless compartment, capable of encapsulating molecules of interest in a DNA-programmed manner. Since the phase separation is driven by the complementarity of DNA sequences at the sticky ends in DNA motifs, this system enables phase separating into multiple phases with possible DNA sequence arrangements, resulting more independent phases than classical binary systems^[20]^. Their phase behaviors in solution^[21,22]^, control of its dynamics^[19,23,24]^, and applications such as protein localization^[25]^ has been studied or/and achieved. Recently, the condensates of those nanomotifs have also been studied with lipid bilayers: for example, DNA nanomotif’s condensates are recruited via electrostatic interaction with cationic lipids doped in lipid monolayers in water-in-oil droplets, forming phase separation on lipid layers^[26]^. In addition, the preexisting motif’s droplets are tagged with membranes via cholesterol modification and their contacts with supported lipid bilayers (SLBs) are photo-controlled, eventually to induce budding in GUVs^[27]^. The large unilamellar vesicles (LUVs) has been encapsulated into the DNA droplet for a control in vesicle transports^[28]^. The above-listed works focuses primarily on the contacts between preexisting DNA nanomotif’s droplet and membranes. In contrast, the behavior of membrane-anchored DNA nanomotifs distributed in two dimensions on lipid bilayers remains unexplored.

In this study, we explore the potential of the DNA nanomotifs for spatially mapping of a GUV surface. Unlike the previous works focusing on the contact between preformed DNA motif’s condensates and lipid bilayers, we studied the phase behavior of two-dimensionally distributed DNA nanomotifs anchored to the lipid bilayers of GUVs via cholesterol modification. When composed of only the motifs with complementary sticky ends, the motifs distribute homogeneously on GUV membranes whereas mixtures of orthogonal two different motifs undergo two-dimensional segregation into gel-like domains with limited lateral mobility in motifs on bilayers. Notably, this organization does not require lipid phase separations, demonstrating that DNA-driven patterning can be achieved independently of membrane phase separation. Our findings provide a foundation for integrating functional membrane domains mapping with phase-separating DNA scaffolds, offering new avenues for designing spatially ordered membranes in synthetic cells.

## Results and Discussion

### Design and characterization of DNA nanomotifs in solution

The DNA nanomotifs investigated in this study consist of four-armed star constructs, as depicted in **Fig. 1A** (sequences are provided in **Supplementary Tables S1 and S2**). Each arm consists of 16 base pairs of “stem” region, with homologous sticky end base-paring with each other at the extremity of each arm. A flexible TT junction at the center of the structure reduces steric constraints and enables the formation of a four-arm configuration. Formation of designed star constructs requires four single-stranded DNAs (ssDNAs), and one of the strands is modified with cholesterol for anchoring to the lipid bilayer via hydrophobic interaction. This design was based on the previous studies of four-armed DNA nano-motifs^[19,23]^, whose structural principles were maintained except for the modification with cholesterol. The assembly of these motifs involved thermal annealing of a cocktail of short complementary oligonucleotides, following established protocols^[19]^. Through hybridization of complementary sequences, four ssDNAs self-assemble into a four-armed star structure (**Fig. 1A**). The choice of sticky-end sequences was based on the previous studies of DNA nanomotifs in solution^[19,23]^. Note that we excluded motifs featuring TATA sticky-ends, as these were previously shown to fail condensate formation^[20]^. Instead, we worked on other sequences of 4nt and 6nt sticky-end sequences which have lower transition temperatures between droplet-like state to gel state (10 □C and 20 □C for 3 arm Y-shape motifs with 4nt and 6nt sticky ends in 20mM Tris pH 8 containing 350 mM NaCl, respectively^[19]^) when it forms condensates in solution. The resulting motifs, S-, and X-motifs studied in this work, are distinguished by their unique sticky-end sequences as described in **Fig. 1A, *top right***.

**Figure 1:**
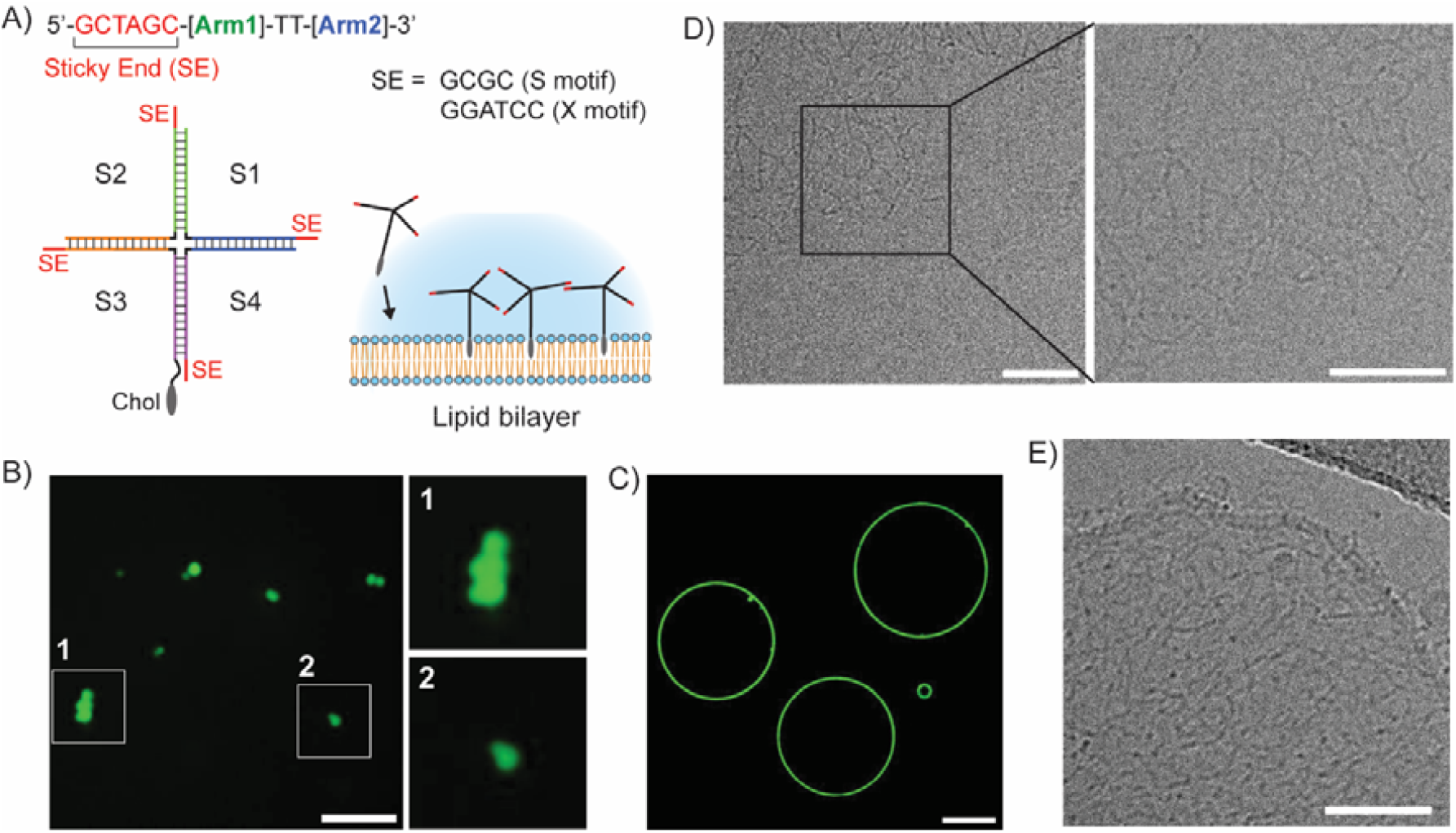
(A) Schematic representations of a DNA motif modified with cholesterol. Sticky ends at 5’, and two sequences for the arms with TT junction in between. The arms are formed by hybridization of complementary sequences of the others. Sequences of sticky ends used in our study are listed, *top right*. Full sequences of all motifs used in this study is available in **Supplementary Table S1 and S2.** (B) A fluorescence microscopy image of S-motif labelled with 10% of Atto488 modified S1 oligo at 1 μM in 20 mM Tris pH 8 containing 100 mM NaCl. Scale bar: 10 μm. The numbered square regions are zoomed on the left (size of images: 10 x 10 μm). (C) A fluorescence microscopy image of S-motif mixed with EggPC GUVs, at 0.75 μM of S-motif in 15 mM Tris pH 8 containing 75 mM NaCl. Scale bar: 10 μm. Cryo-electron microscopy images of (D) S-motifs in solution (top left of the image), and (E) S-motifs anchored to the EggPC LUVs in 20mM Tris pH8 containing 100mM NaCl. Scale bars are 50 nm.

We first examined the behavior of the four-arm nanomotifs in solution. **Figure 1B** presents a fluorescence microscopy image of S-motifs (sequences shown in Materials and Methods), incorporating 10% Atto488-labeled oligonucleotides at 1 µM of DNA (per oligonucleotide) in 20 mM Tris pH 8 with 100 mM NaCl. At room temperature ca. 18 □C, gel-like aggregates with anisotropic shape have been observed. This observation is consistent with prior reports on 4 arm star-shaped motifs^[23,29]^. Initially, we have formed the nanomotifs at high salt buffers, at 20 mM Tris pH 8 with 350 mM NaCl, following the same conditions as previous studies (**see Supp. Fig. S1**). By varying the DNA and NaCl concentrations we found that the S-motif undergoes phase separation in solution to form condensates at DNA concentrations exceeding ∼100 nM (per oligonucleotide) and NaCl concentrations at above 50 mM (**Supp. Fig. S1**). Similar experiments have been performed for X-motifs to examine the difference behaviours between motifs with 4nt sticky end and 6nt sticky end. The X-motif with 6nt sticky end exhibits phase separation under comparable conditions at room temperature as S-motif with 4nt sticky end (**Supp. Fig. S2**).

### Membrane anchoring of DNA nanomotifs

The anchoring and the phase behavior of DNA motifs on two-dimensional GUV surfaces were investigated using fluorescence confocal microscopy. At room temperature, the motifs formed was added to the exterior of giant unilamellar vesicles (GUVs) composed of Egg PC lipids, while ensuring osmotic equilibrium between the GUV interior and exterior. The DNA motifs spontaneously anchored to the lipid bilayers, and the GUV surface becomes enriched with DNA motifs as shown in **Fig. 1C**. Mixing the GUVs with DNA motifs without cholesterol modification shows hardly any anchoring of motifs to GUV membranes (see **Supp. Fig. S3**), demonstrating that hydrophobic insertion of cholesterol is required for anchoring. The structural integrity of the motifs and their membrane association were further confirmed by cryo-electron microscopy. **Figure 1D** and **1E** depict S-motifs in solution and the motifs anchored to Egg PC large unilamellar vesicles (LUVs), respectively. The DNA motifs studied in high salt buffers in previous works^[19,23]^ maintain their proper formation even in relatively low salt conditions as shown in **Fig. 1D** (with 100 mM NaCl). In our experiments, upon mixing motifs with GUVs, motif’s solution can get diluted in 0.75 folds, resulting in final NaCl concentration at 75 mM. We have thus performed the cryo-EM for the samples prepared at 50 mM NaCl and have observed that the structures can still be maintained at 50mM (see **Suppl. Fig. S4**). The cryo-EM images of motifs anchored to the LUVs, in **Fig. 1E** and **Supp. Fig. S4**, show that the motifs anchored to membranes assemble into network-like structures of star-shaped objects, via interactions between the sticky ends at extremities of motif’s arms. This tendency is consistent with the motif’s design and is observed in both solution and on membranes when suspended.

Membrane-anchored S-motifs via cholesterol tag, unlike their behavior in solution, do not undergo phase separation on the two-dimensional (2D) lipid bilayer but instead distribute homogeneously across the GUV surfaces (**Fig. 1C**). This result was consistent over a wide range of NaCl (75mM to 350mM) and DNA (0.075 to 0.75 µM) concentration (see **Supp. Fig. S5**). Interestingly, we observed that when the orthogonal motifs were mixed, these motifs undergo phase separation on GUV membranes. The mixture of S- and X-motifs in 1 to 1 ratio has been prepared by simply mixing two solutions, maintaining the total DNA concentration while decreasing the concentration of each motif. The S- and X-motif mixture forms coexisting gels non-interacting in solution as shown in **Fig. 2A** (S- and X-motifs labeled with Atto488 and Atto550, respectively). When the GUVs are introduced into the solution, the motifs anchored to the Egg PC membranes of GUVs exhibits the segregation as shown in **Fig. 2D**. Indeed, a time-resolved observation presented in **Fig. 2B and 2C** show gradual anchoring of both motifs to the lipid bilayers. First, the strong enrichment of two motifs segregating at the contact of two GUVs happens, when two GUVs are close by. This is due to the interaction between motifs through sticky-end, which is not limited within the 2D lateral membrane surfaces, but can be three-dimensional forming networks between two GUV surfaces. After, the motifs exhibit phase separation on GUV surfaces within around 10-15 minutes, reaching saturation approximately at around 40 minutes at room temperature (**Fig. 2B**). After phase separation occurs, the shape and behaviours of domains do not change significantly. The sample examined in this study has at least incubated 30 minutes before start imaging for the analysis of phase separation. As shown in **Fig. 2D and 2E**, the anchored S- and X-motifs segregated into distinct domains, while the lipids (Egg PC, stained with 1% Cy5-DSPE) remained homogeneously distributed. Rhodamine DHPE, a fluorescently labelled lipid used as reporter for lipid phase separation^[30]^ were also homogeneously distributed in the presence of motif’s phase separation (see **Supp. Fig. S6**), suggesting that the observed phase separation of DNA motifs is not induced by the phase separation of lipid bilayers. The obtained results resemble the results shown in previous work^[26]^, where preformed droplets of star-shaped DNA nano-motifs are bound to cationic-lipid doped lipid monolayer via electrostatic interaction and similar gel-like phase-separating domains has been observed in a mixture of two orthogonal motifs. Importantly, cholesterol enables the motif to integrate more deeply into bilayers compared to electrostatic interactions with cationic lipids; nevertheless, similar macroscopic assemblies of DNA motifs are ultimately observed.

**Figure 2:**
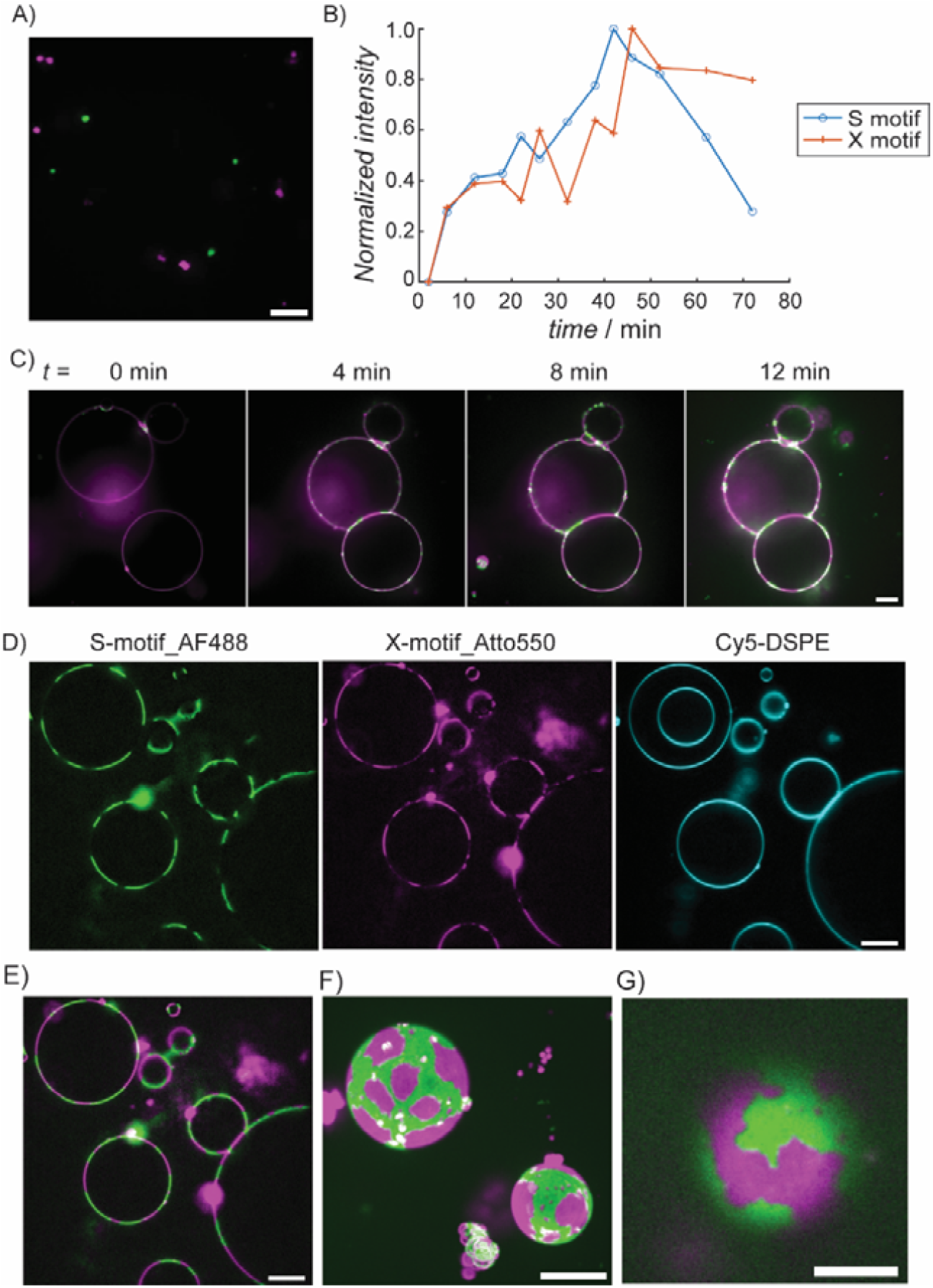
(A) A fluorescence microscopy image of S-motif and X-motif, labelled with 10% of Atto488 and Atto550 respectively, at 1 μM (0.5 μM S-motif, 0.5 μM X-motif) of DNA in 20 mM Tris pH 8 containing 100 mM NaCl. S-motif and X-motifs are presented as green and magenta, respectively. Scale bar: 10 μm. (B, C) Time lapse observation of motif’s insertion to GUV surfaces at 0.75 μM of motifs (S:X = 1:1) in 15 mM Tris pH 8 containing 75 mM NaCl. B-an intensity of motifs across membranes and its evolution over time. C-Time lapse images of GUVs incubated with S- and X-motifs. (D) Fluorescence microscopy images of S- and X-motifs mixed with GUVs stained with 1% of Cy5-DSPE. The sample consists of 0.75 μM of motifs (S:X = 1:1) with 15mM Tris buffer at pH 8 containing 75 mM NaCl. Left: S-motifs, Middle: X-motifs, and Right: Cy5-DSPE. S-motif, X-motifs and Cy5-DSPE are presented as green, magenta, and cyan, respectively. Scale bar: 10 μm. (E) An overlaid image Fig. 2D left (S-motif) and middle (X-motif). Scale bar: 10 μm. (F) Reconstitution of 3D structures from 2D images at different cross-section in z-axis for a half sphere of GUVs. The images at different focal height are projected by Max Intensity in Image J. S-motif and X-motifs are presented as green and magenta, respectively. Scale bar: 10 μm. (G) A zoomed image of a bottom of phase-separated GUV. Scale bar: 5 μm.

### Characterizing the physical state of DNA domains formed on lipid bilayers

Regarding the physical state of domains, three-dimensional reconstructions of GUVs decorated with two orthogonal DNA motifs (**Fig. 2F and 2G**) show anisotropic shape of domains with non-smooth phase boundaries, adopting a solid or gel-like state. This was further confirmed by performing fluorescence recovery after photobleaching (FRAP) to see the mobility of motifs within and across domains. Representative FRAP recovery data presented in **Fig. 3A *top*** shows partial photobleaching of a domain region, while **Fig. 3A *bottom*** illustrates full photobleaching of an isolated domain composed of S-motifs on GUV surfaces. For FRAP, samples have been incubated one hour to characterize domains at the steady state, and GUV instability has been minimized by a strong density contrast between the internal sucrose solution and the external buffer containing NaCl, which let GUV sedimented and more immobile. However, imaging the top surface of GUVs over extended periods proved challenging due to focal drift. The domains located at the GUV bottom has thus been analysed. The limited sampling size reflects both the difficulty in maintaining a stable focal plane for prolonged observations and the rarity of isolated domain formation at the GUV bottom. **Figures 3B and 3C** display the normalized recovery curves averaged across repeated experiments. Partial photobleaching of a domain revealed a slow fluorescence recovery (**Fig. 3B**), persisting even after three minutes of acquisition. The half-times for recovery ( *t*_*1/2*_ ) and mobile fractions ( *f*_*m*_) were as follows: for S-motifs (*n* =5), *t*_*1/2*_ 82 ± 67s and *f*_*m*_ = 0.40 ± 0.26; for X-motifs (*n* = 6), *t*_*1/2*_ =69 ± 59s and *f*_*m*_ = 0.29 ± 0.16 (mean ± SD). Full bleaching of an entire domain also resulted in slow fluorescence recovery (**Fig. 3C**), having a plateau at the end, with values for S-motifs (*n* = 4): *t*_*1/2*_ = 39 ± 24 s and *f*_*m*_ = 0.13 ± 0.04 ; and for X-motifs (*n* = 4 ): *t*_*1/2*_ = 55 ± 20 s and *f*_*m*_ = 0.10 ± 0.07(mean ± SD). Characteristic timescales and mobile fractions were obtained from the FRAP curves fitted with a single exponential function (details in Materials and Methods and in **Supp. Fig. S7**). Note that out-of-focus frames were excluded from the analysis, that varies the duration of acquisition time and the amount of data for the obtained mean values close to the end of acquisition. The observed fluorescence recovery may arise from several mechanisms, including lateral diffusion of motifs across domains, diffusion within domains, and exchange with the surrounding solution. In experiments involving partial and full bleaching of domains, both S- and X-motif domains showed half-times with high variability. The high variability in half-times suggests the absence of a single characteristic timescale, indicating high heterogeneity across GUVs. Full domain photobleaching revealed a similar recovery timescale but with a reduced mobile fraction compared to the partial bleaching. The higher mobile fraction observed in partial bleaching, compared to full bleaching, suggests possible motif exchange within a domain. Overall, the results demonstrate gel-like behavior, characterized by slow recovery rates and low mobile fractions. This contrasts with the rapid diffusion of lipids in fluid lipid membranes, which typically exhibit mobile fractions of 70–90% and half-times of a few seconds^[31]^. This suggests that the interconnected networks of DNA motifs on membranes exhibit low reconfigurability at room temperature. The variability in results may reflect sample heterogeneity, including differences in phase distribution, the number of anchored motifs in GUVs, and the surrounding DNA nanomotif condensates, where motif’s exchange with the solution may not be negligible. Addressing these limitations will be essential for developing more robust and controlled systems.

**Figure 3:**
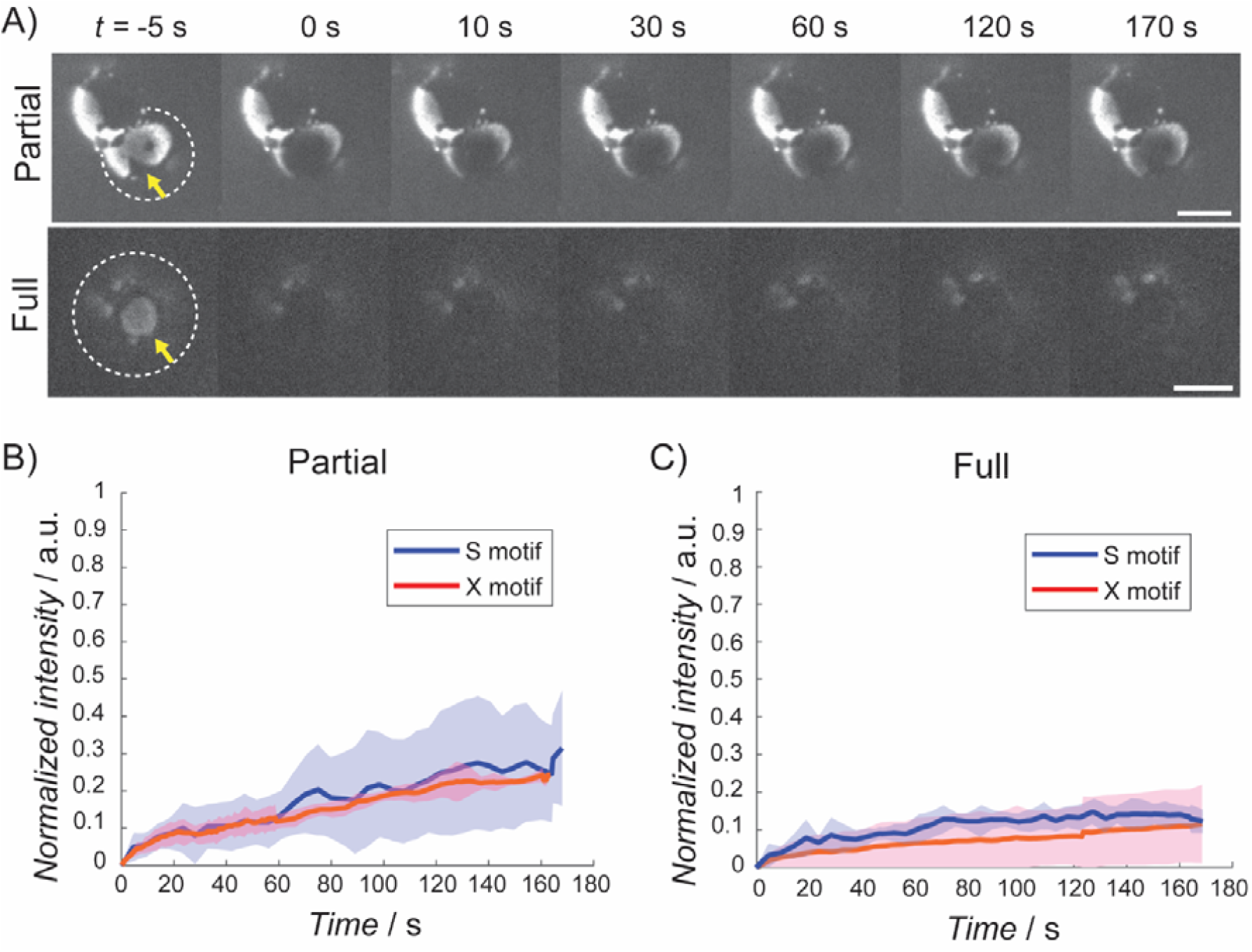
Temporal dynamics and recovery of fluorescence after photobleaching (FRAP) of S- and X-motifs anchored to lipid bilayers of GUVs at 0.75 μM of total motif’s concentration (S:X = 1:1 ratio) in 15 mM Tris pH 8 containing 75 mM NaCl. (A) Representative time-lapse images of (*bottom*) a partially photobleached domain of S-motifs, and (*top*) a fully photobleached isolated domain showing fluorescence recovery at the indicated time points. Scale bars: 5 µm. Images were captured at the bottom of GUVs, with dashed white lines indicating the GUV bottom. Yellow arrows highlight the photobleached domain (full bleaching) and region (partial bleaching). Normalized fluorescence intensity over time for S-motif (blue) and X-motif (red) (B) in a partial photo bleaching of a domain and (C) in a full bleaching of an isolated domain. The solid line indicates the averaged values with the standard deviations depicted as bands. Sample sizes: S Partial (n = 5), X Partial (n = 6), S Full (n = 4), X Full (n = 4).

### Phase behaviors of membrane-anchored DNA nanomotifs on membranes

The behavior of membrane anchored DNA nanomotifs can be governed by a balance between three key interactions. As DNA is a negatively charged polyelectrolyte, repulsive force between motifs depends on the ionic strength of the solution, specifically, the NaCl concentration in this system. In contrast, complementary sticky ends increase attraction between homologous motifs. Additionally, hydrophobicity of cholesterol enables anchoring to lipid bilayers. Together, these three weak interactions, *i*.*e*., electrostatic repulsion, sticky end attraction, and hydrophobic anchoring, would either compete or assist, that govern the phase behaviors of the motifs. In this study, we have investigated how the change in electrostatic interactions by varying NaCl concentration differs the anchoring of the motifs to membranes, as well as the phase behaviors of motifs on membranes. **Figure 4A** presents a phase diagram displaying the behaviors of S- and X-motifs on GUV surfaces in varied DNA and NaCl concentrations, supplemented with representative fluorescence microscopy images corresponding to each condition in the phase diagram (unzoomed images at each condition are displayed in **Supp. Fig. S8**). At DNA concentration above 100 nM and NaCl concentration at 75 mM, two orthogonal motifs segregate on the GUV surfaces, as shown in **Fig. 2D**. Similarly, at 150 mM NaCl, motif segregation occurs but requires relatively higher DNA concentrations compared to the low-salt condition (**Fig. 4A *left*)**. Under high-salt condition (300 mM NaCl), commonly used in previous studies of similar DNA stars ^[19,23]^, fewer motifs anchored to the membranes. Instead, they localize predominantly at the contact sites between two GUVs (**Fig. 4A *right***), while gel-like condensates persist in bulk solution. Increasing NaCl concentration reduces enrichment of motifs on GUV membranes. This behavior is likely driven by electrostatic screening, which weakens repulsion between negatively charged DNA motifs. Indeed, the higher ionic strength can enhance the sticky end interactions, potentially modulating the concentration of motifs in diluted phase, or favouring the associated condensate states in bulk over membrane anchoring through cholesterol’s hydrophobicity. The balance between sticky end interactions, electrostatic interactions between motifs, and hydrophobicity of cholesterol, is critical for the anchoring and subsequent phase separation of motifs on GUVs. Similarly, to the behaviors of motifs in high salt buffer, the enrichment of anchored motifs at the contact of two GUVs is also observed at low DNA concentrations under low-salt conditions (75 mM NaCl), suggesting that the limited numbers of motifs on membranes preferentially localize at the contact between two GUVs (see **Supp. Fig. S8**). This may result from a preferential local enhancement of sticky end-driven interactions, occurring both laterally within bilayers and vertically between bilayers during vesicle contact.

**Figure 4:**
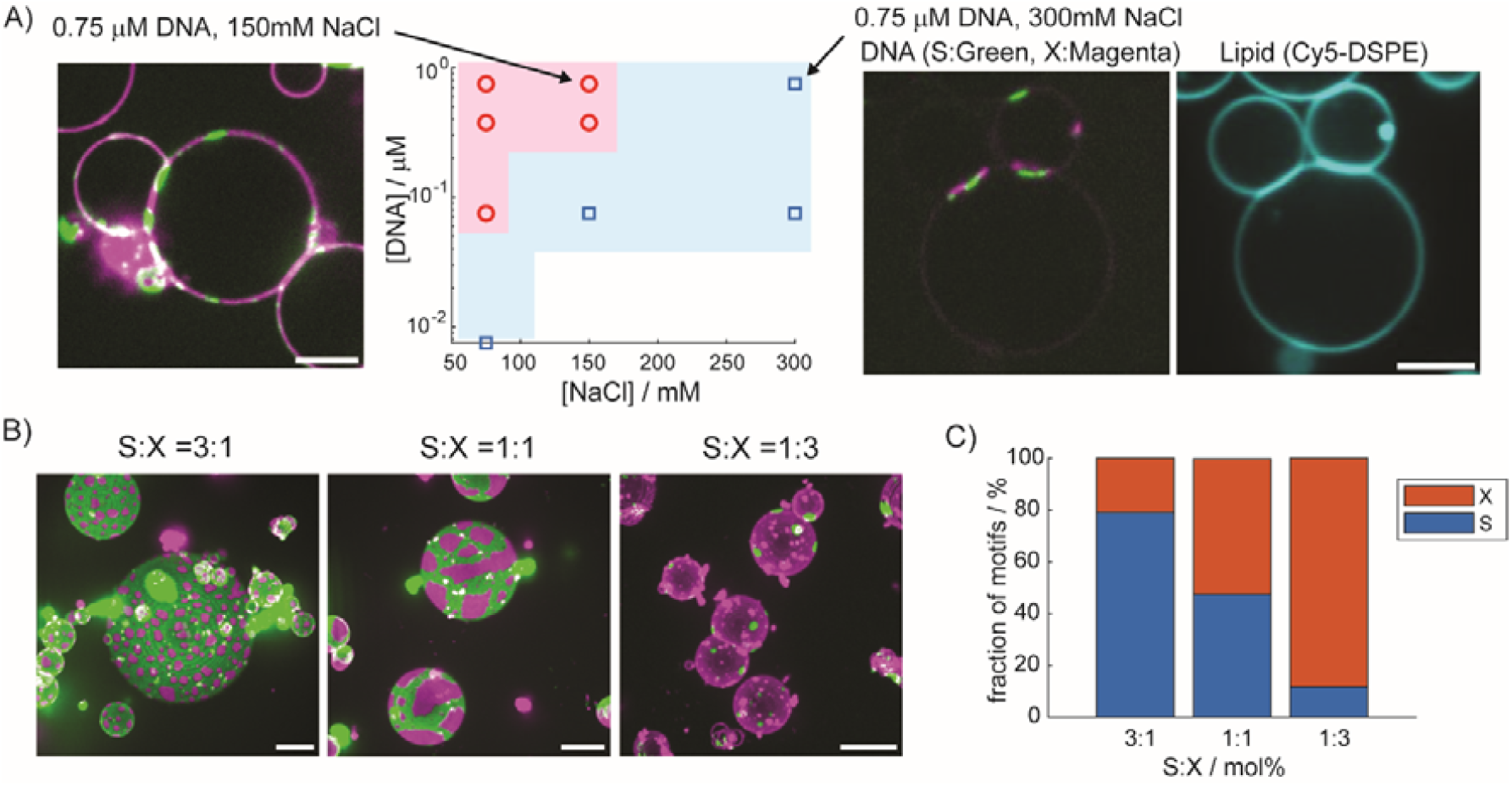
(A) A phase diagram presenting the behaviors of a mixture of S- and X-motifs (1:1 ratio) on GUVs as a function of DNA and NaCl concentration. The red circle highlights the distribution of both S- and X-motifs within membranes exhibiting segregation. The square marks the enrichment of these motifs at the two GUV contact sites, while the free membrane regions show no such motif distribution. The representative fluorescence microscopy images of S- and X-motifs at both conditions are shown in left and right of the phase diagram, respectively. S- and X-motifs are both visualized as green and magenta, respectively. All scale bars are 10 μm. B) Fluorescence microscopy images of S- and X-motifs anchored to GUVs in different stoichiometry of two motifs. The images at different cross-section of GUVs are projected with Max Intensity in ImageJ. All scale bars are 10 μm. (C) The bar graph shows the fraction of S- and X-motifs on GUV surfaces incubated with different ratio in S- and X-motifs.

In the mixture of orthogonal motifs on lipid bilayer, the role of stoichiometry in fraction of phases composed of each motif on membranes has been studied by adjusting the composition of motifs in solution. **Figure 4B** presents fluorescence microscopy images of GUVs incubated with S- and X-motif cocktails at different ratios (S:X = 3:1, 1:1, and 1:3 in molar ratio). **Figure 4C** further quantifies the fraction of the GUV interface covered by each motif at these varied ratios from analysis of GUVs (number of experiments: *N* = 2, number of vesicles: *Nv* = 10 for each condition). The area covered by each motif on the GUV surface correlates with its relative concentration in solution: the majority phase comprises the more abundant motif, while the minority motif forms distinct domains. Interestingly, when S- and X-motifs are present in equal proportions, the X-motif exhibits a slightly greater tendency to cover the GUV surface. This discrepancy may arise from differences in the strength of interactions between homologous motifs, which are influenced by the length and sequence of the sticky ends, S-motifs feature a 4-nucleotide (GCGC) sticky end, whereas X-motifs have a longer 6-nucleotide (GGATCC) sequence. The stronger attraction between X-motifs might enhance their recruitment and anchoring to the membrane in corrective manner, resulting in their increased surface coverage compared to S-motifs.

## Conclusion

We studied the phase behavior and the dynamics of membrane anchored DNA nanomotifs on lipid bilayers of giant unilamellar vesicles. Our results demonstrate that four-armed star-shaped DNA nanomotifs, functionalized with a cholesterol moiety on one arm, spontaneously anchor to lipid bilayers. When the DNA motifs possess complementary sticky end sequences, they interact each other and distribute homogeneously along lipid bilayers. While a mixture of two orthogonal motifs with different sticky ends undergoes phase separation, forming a gel-like domain with slow dynamics in motif’s exchanges inside the domain as well as across isolated domains. Motif’s enrichment on membranes occurs only at relatively low salt concentrations, reflecting a balance between electrostatic interactions among motifs and hydrophobicity of cholesterol. The phase separation observed in mixtures of two orthogonal motifs anchored to bilayers via cholesterol-tag produce similar patterns observed in the previous studied system, where preexisting DNA motif’s droplets are bound to membranes doped with cationic lipids via electrostatic interactions^[26]^. Importantly, lateral DNA phase separation observed in this study through cholesterol-anchoring does not require specific lipids in underlying lipid composition, providing us more flexibility in the system. As a future perspective, given the possibility of DNA sequences at the sticky ends, the studied system may be extended to multi-phase separation on membranes to overcome the limitations of classical binary phase separation systems, typically observed on lipid phase separations. Coupling the inherent ability of the star-shaped DNA nanomotifs in recruitment of the molecules of interests as shown in previous works^[19,25]^, and integrating the other strategy (e.g., strand-displacements^[23]^, or cross-bridging motifs to ligase the domains^[19]^) would enable us to spatially and temporally control the arrangement of membrane associated proteins and other molecules in DNA encoded manner. In analogy with previous applications of DNA (or RNA) nano stars for the localization of specific components and integration to *in vivo* system^[25,28,32,33]^, studied two-dimensional domains of DNA nanomotifs on lipid bilayers may offer a promising approach to spatially organize and regulate membrane components in cell surfaces.

## Materials and Methods

### Chemicals

Acetone, Chloroform, Isopropanol, Sodium Chloride, Potassium Hydroxide, Trizma base, Sucrose, 1-(6-((2-((((R)-2,3-bis(stearoyloxy)propoxy)(hydroxy)phosphoryl)oxy)ethyl)imino)-6-hydroxyhexyl)-3,3-dimethyl-2-(5-(1,3,3-trimethylindolin-2-ylidene)penta-1,3-dien-1-yl)-3H-indol-1-ium (Cy5-DSPE) and L-α-phosphatidylcholine (Egg, Chicken) (Egg PC) are all purchased from Sigma Aldrich Ltd.

### Formation of DNA nano-motifs with four arms

The oligonucleotides were purchased from IDT DNA (Leuven, Belgium) and were prepared by standard desalting, except for the fluorophore-labeled and cholesterol modified strands, which were purified to high-performance liquid chromatography (HPLC) grade. Sequences of the oligonucleotides are provided in **Supplementary Tables S1 and S2**. The oligonucleotides were dissolved in ultrapure water (18 MΩ · cm in resistance) at 100 µM or 1 mM concentrations and stored at −20°C until use. Cholesterol-modified oligos were heated to 60° C for 5 minutes before the use, following the protocol described in previous work^[34]^. The desired amount of DNA strands was mixed in a test tube with Tris buffer and NaCl solution to be finally at 20 mM Tris buffer at pH 8 with desired concentration of NaCl. The test tube was heated at 95°C for 3 min and then gradually cooled down to 25°C at a rate of −1°C/min using a thermal cycler (T100 Thermal Cycler, BIO-RAD), as described in previous work^[19]^. The annealed samples were then transferred directly to the chambers or mixed with giant unilamellar vesicles in eppendorf tube. Protocol for the passivation of observation chambers was described below.

### Electroformation of Giant Unilamellar Vesicles

GUVs were prepared by the electroformation method using a custom-made protocol adapted from the historical paper^[35]^. Egg PC in chloroform purchased were diluted by chloroform to a total lipid concentration of 1 mg/mL. Cy5-DSPE was added to the lipid mixture to be 1 mol% in total lipid composition in order to visualize the lipid bilayer of GUVs, respectively. 10 μL of prepared lipid mixture was spread on the conductive side of ITO-coated glass slides (surface resistivity: 15-25 Ω/sq). A lipid dry film immediately formed on the substrate and the film was further dried under vacuum for more than 30 minutes. A chamber made by the two lipid-deposited ITO slides and roughly 1 mm-thick PDMS spacer was then filled with a desired concentration of Sucrose solution, which osmolarity is equilibrated with the external solutions containing buffer, salts and oligonucleotides. Osmolarity was measured by Vapor Pressure Osmometer (VAPRO, ELITechGroup), Freezing point Osmometer (Loser). The chamber was sealed with a PDMS plug, and 1.1 Vp-p AC voltage at 10 Hz was applied for 3 hours, followed by the detachment with 1.1 Vp-p at 5 Hz for 30 minutes using function generator (TG315, Aim-TTi). The resulting GUVs were harvested by pipettes to a test tube and kept at 4 ºC and used for experiments within one week.

### Passivation of the observation chamber

For the passivation of glass substrates, we have used a protocol applied in previous work^[36]^. Glasses and coverslips, measuring 22 mm by 40 mm and 20 mm by 20 mm, respectively, with a thickness No 1,5 were purchased from VWR. The cleaning and coating procedure of the glasses are as follows: Cover glasses were washed with for 20 minutes under sonication. It is followed by rinsing with distilled water and then washing with 1M KOH for 20 minutes under sonication. After rinsing with distilled water and wash with distilled water for 20 minutes of sonication, the cover glasses are dried with nitrogen gas. The surface of glasses was then siliconized with Sigmacote, and then were rinsed with Isopropanol, and were dried with nitrogen gas, and were kept until the use. To assemble the chamber, silicone double sided tapes (SecureSeal− Adhesive Sheets, Grace Bio-Lab) acting as spacers are sandwiched between a cover glass and a slide glass, or cover glass bottom with custom PDMS wells. The chamber is then incubated with 0.5% pluronic acid (PF127) for 15 minutes. The chamber is then rinsed with the 20mM Tris pH 8 buffer few times and then followed by the addition of samples. For long time acquisition, the chambers with two glasses sealed with glue are used, or for custom PDMS wells, additional cover slide is placed on custom PDMS well to avoid evaporation.

### Addition of DNA nanomotifs to Giant Unilamellar Vesicles

DNA nanomotifs, observation chambers and GUVs were prepared as described above. DNA nanomotifs were mixed with GUVs suspensions directly in the observation chamber. For all experiments, DNA nanomotifs were added to the GUVs suspension at a fixed volume ratio of 3:1 (DNA nanomotifs : GUVs), with a final total volume of 60 µL per chamber. Unless otherwise stated, mixtures of S- and X-motifs were prepared at an equimolar ratio (1:1). For experiments investigating the effect of stoichiometry, the relative proportions of S- and X-motifs were varied while maintaining the total DNA concentration constant. After mixing, samples were gently mixed by pipettes and incubated at room temperature prior to imaging. All observations were performed under conditions of osmotic equilibrium between the internal and external solutions.

### Fluorescence microscopy imaging

Inverted Eclipse Ti-E (Nikon) confocal microscope equipped with Spinning disk CSU-X1 (Yokogawa) integrated in Metamorph software was used. Images are taken by Prime 95B (Photometrics) and are further analyzed using imageJ and Matlab softwares described just below:

#### Membrane Enrichment

Time-lapse image sequences were analyzed to quantify fluorescence enrichment at the GUV membrane. For each vesicle: A circular ROI was drawn along the membrane. A band (thickness ∼5–6 pixels) was generated using the *Make Band* function in Fiji to capture membrane-associated signal. Mean fluorescence intensity within the membrane band was measured over time. Background subtraction was applied using a region devoid of fluorescent signal.

#### Stoichiometry Analysis of DNA Motifs

To quantify the relative distribution of DNA motifs (S and X) along the GUV membrane, a line-based analysis was performed. Images were first corrected for background and thresholded to generate binary masks for each channel. A line ROI was drawn along the membrane and straightened using the *Straighten* function in Fiji. Fluorescence intensity profiles for both channels were extracted using *Plot Profile*, yielding pixel-wise intensity values along the membrane contour. For each pixel: intensities from both channels were compared, the pixel was assigned to the motif with a higher intensity. The percentage of membrane area occupied by each motif was calculated as:

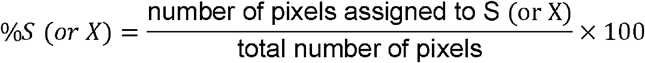

#### Fluorescence Recovery after Photo-bleaching

Image sequences for FRAP were analyzed using ImageJ: A circular ROI was drawn to encompass the region of fluorescence recovery. A second circular ROI was selected to measure background fluorescence. For each ROI, the mean fluorescence intensity was measured over time. The obtained fluorescence intensity profiles have been normalized, and the normalized data were fitted with a single exponential function 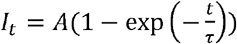 to obtain the half-times for recovery (*t*_*1/2*_ = *τ* log (2)) and mobile fractions (*f*_*m*_=*A*).

### Cryo-electron microscopy

Lipid mixture (Egg PC 100%) dissolved in chloroform was quickly dried under nitrogen. Obtained lipid dried films are further dried under vacuum for more than 30 min. Lipids are resuspended in 20mM Tris pH 8 buffer containing desired amount of NaCl by vortex to obtain various size of LUVs. LUVs suspension was diluted to 0.12 mg/mL and DNA motifs were added to be a target final concentration. Mixture was incubated for more than 30 minutes, and 4 μL of solution was deposited on a glow discharged lacey carbon electron microscopy grid (Agar Scientific, United Kingdom). Most of the solution was blotted away from the grid to leave a thin (<100 nm) film of aqueous solution. The blotting was carried out on the opposite side from the liquid drop and plunge-frozen in liquid ethane at −181 ºC using an automated freeze plunging apparatus (EM-GP2, Leica, Germany). The samples were kept in liquid nitrogen and imaged using a Glacios microscope (Thermo Fisher, USA) operating at 200 kV with a falcon IVi camera and in low dose mode.

## Acknowledgement

We deeply acknowledge Daniel Lévy, Patricia Bassereau, Simli Dey, Feng-Ching Tsai, Aurélie Bertin, Manuela Dezi, Julien Maufront, John Manzi, Benoit Schaeffer-Dancoisne, Uriel Fuentes Beltran, and Carine Vias for the experimental support or/and discussion. We thank Pulkit Aditya and Johanna Gerstenecker for the support in image analysis. We thank Vincent Fraisier, Olivier Leroy, Chloe Guedj, Amelie Sarrazin and the other members of PICT-IBiSA in Institut Curie for the support on optical microscopy imaging.

## References

[1] K. P. Adamala, M. Dogterom, Y. Elani, P. Schwille, M. Takinoue, T. Y. D. Tang, “Present and future of synthetic cell development” Nat. Rev. Mol. Cell Biol. 2024, 25, 162–167.

[2] S. Jeong, H. T. Nguyen, C. H. Kim, M. N. Ly, K. Shin, “Toward Artificial Cells□: Novel Advances in Energy Conversion and Cellular Motility” Adv. Funct. Mater 2020, 1907182, 1–26.

[3] C. D. Crowe, C. D. Keating, “Liquid–liquid phase separation in artificial cells” Interface Focus 2018, 8, 20180032.

[4] M. Yanagisawa, S. Nigorikawa, T. Sakaue, K. Fujiwara, M. Tokita, “Multiple patterns of polymer gels in microspheres due to the interplay among phase separation, wetting, and gelation” Proc. Natl. Acad. Sci. U. S. A. 2014, 111, 15894–15899.

[5] R. Sakamoto, M. P. Murrell, “Mechanical power is maximized during contractile ringlike formation in a biomimetic dividing cell model” Nat. Commun. 2024, 15, 9731.

[6] R. Harris, N. Berman, A. Lampel, “Coacervates as enzymatic microreactors” Chem. Soc. Rev. 2025, 54, 4183–4199.

[7] G. L. Nicolson, G. Ferreira de Mattos, “The Fluid–Mosaic model of cell membranes: A brief introduction, historical features, some general principles, and its adaptation to current information” Biochim. Biophys. Acta Biomembr. 2023, 1865, 184135.

[8] G. L. Nicolson, G. Ferreira de Mattos, “Fifty Years of the Fluid–Mosaic Model of Biomembrane Structure and Organization and Its Importance in Biomedicine with Particular Emphasis on Membrane Lipid Replacement” Biomedicines 2022, 10, 1711.

[9] K. Y. Lee, S. J. Park, K. A. Lee, S. H. Kim, H. Kim, Y. Meroz, L. Mahadevan, K. H. Jung, T. K. Ahn, K. K. Parker, K. Shin, “Photosynthetic artificial organelles sustain and control ATP-dependent reactions in a protocellular system” Nat. Biotechnol. 2018, 36, 530–535.

[10] H. Jia, L. Kai, M. Heymann, D. A. García-Soriano, T. Härtel, P. Schwille, “Light-Induced Printing of Protein Structures on Membranes in Vitro” Nano Lett. 2018, 18, 7133–7140.

[11] T. Hamada, R. Sugimoto, T. Nagasaki, M. Takagi, “Photochemical control of membrane raft organization” Soft Matter 2011, 7, 220–224.

[12] S. L. Veatch, S. L. Keller, “Separation of Liquid Phases in Giant Vesicles of Ternary Mixtures of Phospholipids and Cholesterol” Biophys. J. 2003, 85, 3074–3083.

[13] T. Baumgart, A. T. Hammond, P. Sengupta, S. T. Hess, D. A. Holowka, B. A. Baird, W. W. Webb, “Large-scale fluid/fluid phase separation of proteins and lipids in giant plasma membrane vesicles” Proc. Natl. Acad. Sci. U. S. A. 2007, 104, 3165–3170.

[14] H. Himeno, N. Shimokawa, S. Komura, D. Andelman, T. Hamada, M. Takagi, “Charge-induced phase separation in lipid membranes” Soft Matter 2014, 10, 7959–7967.

[15] J. Li, A. A. Green, H. Yan, C. Fan, “Engineering nucleic acid structures for programmable molecular circuitry and intracellular biocomputation” Nat. Chem. 2017, 9, 1056–1067.

[16] N. Wu, F. Chen, Y. Zhao, X. Yu, J. Wei, Y. Zhao, “Functional and Biomimetic DNA Nanostructures on Lipid Membranes” Langmuir 2018, 34, 14721–14730.

[17] M. Langecker, V. Arnaut, T. G. Martin, J. List, S. Renner, M. Mayer, H. Dietz, F. C. Simmel, “Synthetic Lipid Membrane Channels Formed by Designed DNA Nanostructures” Science 2012, 338, 928–932.

[18] A. Schoenit, E. A. Cavalcanti-Adam, K. Göpfrich, “Functionalization of Cellular Membranes with DNA Nanotechnology” Trends Biotechnol. 2021, 39, 1208–1220.

[19] Y. Sato, T. Sakamoto, M. Takinoue, “Sequence-based engineering of dynamic functions of micrometer-sized DNA droplets” Sci. Adv. 2020, 6, eaba3471.

[20] A. S. Chaderjian, S. Wilken, O. A. Saleh, “Diverse, distinct, and densely packed DNA nanostar droplets” Proc. Natl. Acad. Sci. U. S. A. 2026, 123, e2523462123.

[21] S. Wilken, A. Chaderjian, O. A. Saleh, “Spatial Organization of Phase-Separated DNA Droplets” Phys. Rev. X 2023, 13, 031014.

[22] S. Biffi, R. Cerbino, F. Bomboi, E. M. Paraboschi, R. Asselta, F. Sciortino, T. Bellini, “Phase behavior and critical activated dynamics of limited-valence DNA nanostars” Proc. Natl. Acad. Sci. U. S. A. 2013, 110, 15633–15637.

[23] S. Agarwal, D. Osmanovic, M. Dizani, M. A. Klocke, E. Franco, “Dynamic control of DNA condensation” Nat. Commun. 2024, 15, 1915.

[24] T. Maruyama, J. Gong, M. Takinoue, “Temporally controlled multistep division of DNA droplets for dynamic artificial cells” Nat. Commun. 2024, 15, 7397.

[25] M. Dizani, D. Sorrentino, S. Agarwal, J. M. Stewart, E. Franco, “Protein Recruitment to Dynamic DNA-RNA Host Condensates” J. Am. Chem. Soc. 2024, 146, 29344–29354.

[26] Y. Sato, M. Takinoue, “Capsule-like DNA Hydrogels with Patterns Formed by Lateral Phase Separation of DNA Nanostructures” JACS Au 2022, 2, 159–168.

[27] N. Kaletta, S. Burick, Y. Qudbuddin, P. Schwille, “Designing Tunable DNA Condensates to Control Membrane Budding Transformation in Synthetic Cells” Adv. Sci. 2025, 12, e15510.

[28] D. A. Tanase, L. Malouf, R. Rubio-Sánchez, K. Jain, B. M. Mognetti, L. Di Michele, “Organization and triggered release of liposomes with DNA-based synthetic condensates” bioRxiv. 2025, DOI 10.1101/2025.09.27.678970.

[29] Y. Sato, M. Takinoue, “Sequence-dependent fusion dynamics and physical properties of DNA droplets” Nanoscale Adv. 2023, 5, 1919–1925.

[30] K. Nakazawa, A. Lévrier, S. Rudiuk, A. Yamada, M. Morel, D. Baigl, “Controlled Lipid Domain Positioning and Polarization in Confined Minimal Cell Models” Angew. Chem. Int. Ed. 2025, 64, e202419529.

[31] B. Chauvin, K. Nakazawa, I. Adriaans, M. Geymonat, S. Piatti, B. Hajj, S. Mangenot, A. Bertin, “Septin Architecture Dictates Size-Dependent Diffusion Barriers on biomimetic Membranes” bioRxiv. 2025, DOI 10.1101/2025.10.08.681182.

[32] G. Fabrini, N. Farag, S. P. Nuccio, S. Li, J. M. Stewart, A. A. Tang, R. McCoy, R. M. Owens, P. W. K. Rothemund, E. Franco, M. Di Antonio, L. Di Michele, “Cotranscriptional production of programmable RNA condensates and synthetic organelles” Nat. Nanotechnol. 2024, 19, 1665–1673.

[33] S. Li, Y. Kim, K. Wang, E. J. Payson, A. A. Tang, M. V. Nieto, D. Osmanovic, M. Yang, D. Dilao, A. Bermudez, W. Xiao, M. M. H. Li, N. Y. C. Lin, K. Plath, D. L. Black, E. Franco, “Programmable artificial RNA condensates in mammalian cells” bioRxiv. 2026, DOI 10.64898/2026.01.28.702393.

[34] A. Ohmann, K. Göpfrich, H. Joshi, R. F. Thompson, D. Sobota, N. A. Ranson, A. Aksimentiev, U. F. Keyser, “Controlling aggregation of cholesterol-modified DNA nanostructures” Nucleic Acids Res. 2019, 47, 11441–11451.

[35] M. I. Angelova, D. S. Dimistov, “Liposome electroformation” Faraday Discuss. Chem. Soc. 1986, 81, 303–311.

[36] R. W. Yao, M. K. Rosen, “Advanced surface passivation for high-sensitivity Studies of biomolecular condensates” Proc. Natl. Acad. Sci. U. S. A. 2024, 121, e2403013121.

