## Supplementary Information for "Two-Dimensional Phase Separation of DNA Nanomotifs Anchored to Lipid Bilayers"


**Table of Contents:**


1. Supplementary Tables S1 and S2
2. Supplementary Figures S1-S8

1. **Supplementary Table**

**Table S1:** Sequences of oligonucleotides for S-motifs used in this study.

| | |
|---|---|
| S1_4_4 | GCGCCAGTGAGGACGGAAGTTTGTCGTAGCATCGCACC |
| S2_4_4 | GCGCCAACCACGCCTGTCCATTACTTCCGTCCTCACTG |
| S3_4_4_chol | GCGCCCATGGTCCCAAGTGATTTGGACAGGCGTGGTTG/3'CholTEG/ |
| S4_4_4 | GCGCGGTGCGATGCTACGACTTTCACTTGGGACCATGG |
| Atto488_S1_4_0 | /ATTO488/ CAGTGAGGACGGAAGTTTGTCGTAGCATCGCACC |

**Table S2:** Sequences of oligonucleotides for X-motifs used in this study.

| | |
|---|---|
| X1_4_6 | GGA TCC AAA GGA ACT CTC CGC GTT GAC AAA GCC GAC ACG T |
| X2_4_6 | GGA TCC GCC TCT GTG TCG CAT CTT CGC GGA GAG TTC CTT T |
| X3_4_6_chol | GGA TCC CAG ACG TCA CTC TCC ATT GAT GCG ACA CAG AGG C/3CholTEG/ |
| X4_4_6 | GGA TCC ACG TGT CGG CTT TGT CTT TGG AGA GTG ACG TCT G |
| Atto550_X4_1_0 | /5ATTO550N/AA AGG AAC TCT CCG CGT TGA CAA AGC CGA CAC GT |

### 2. Supplementary Figures

**Figure S1:** Fluorescence microscopy images of S-motif suspended in solution with varied NaCl and DNA concentrations. Scale bars: 10 μm.

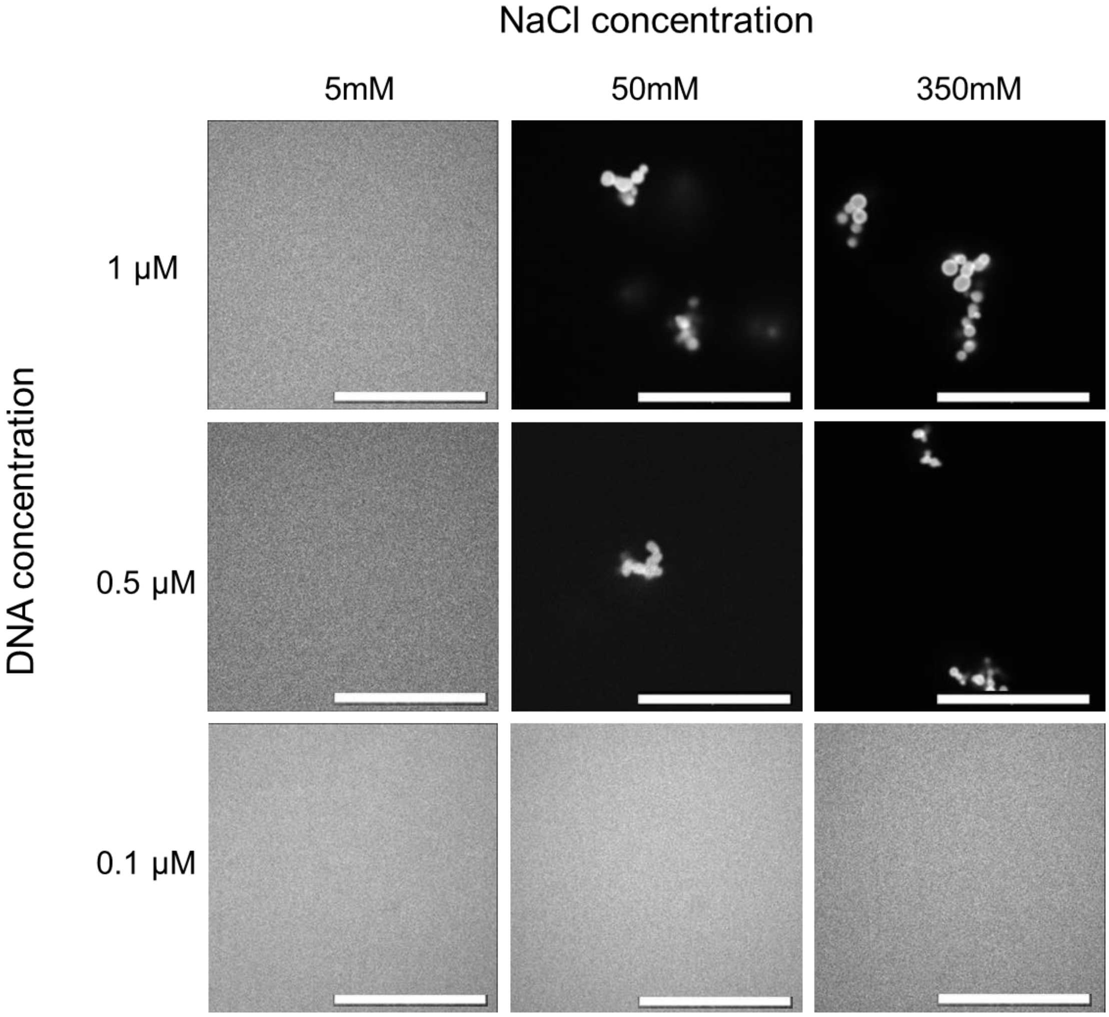

**Figure S2:** Fluorescence microscopy images of X-motif suspended in solution with varied NaCl and DNA concentrations. Scale bars: 10 μm.

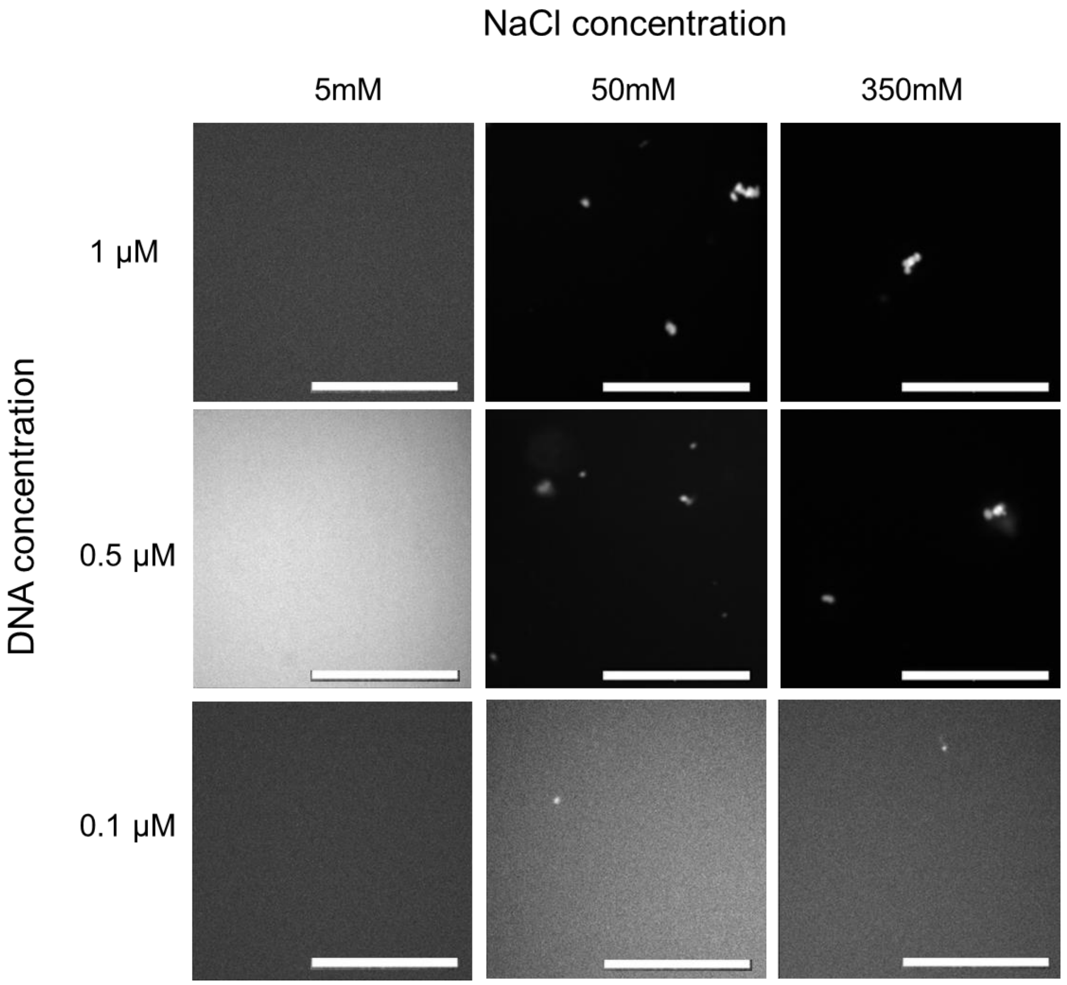

**Figure S3**: Fluorescence microscopy images of S- and X-motifs without cholesterol modifications, mixed with GUVs stained with 1% of Cy5-DSPE. Left: S-motifs, Middle: X-motifs, and Right: Cy5-DSPE. (A) Unzoomed images of the mixture. (B) Zoomed images of GUVs. Contrast was adjusted to enhance the detection of faint motif signals on the membranes. Scale bars: 10 µm.

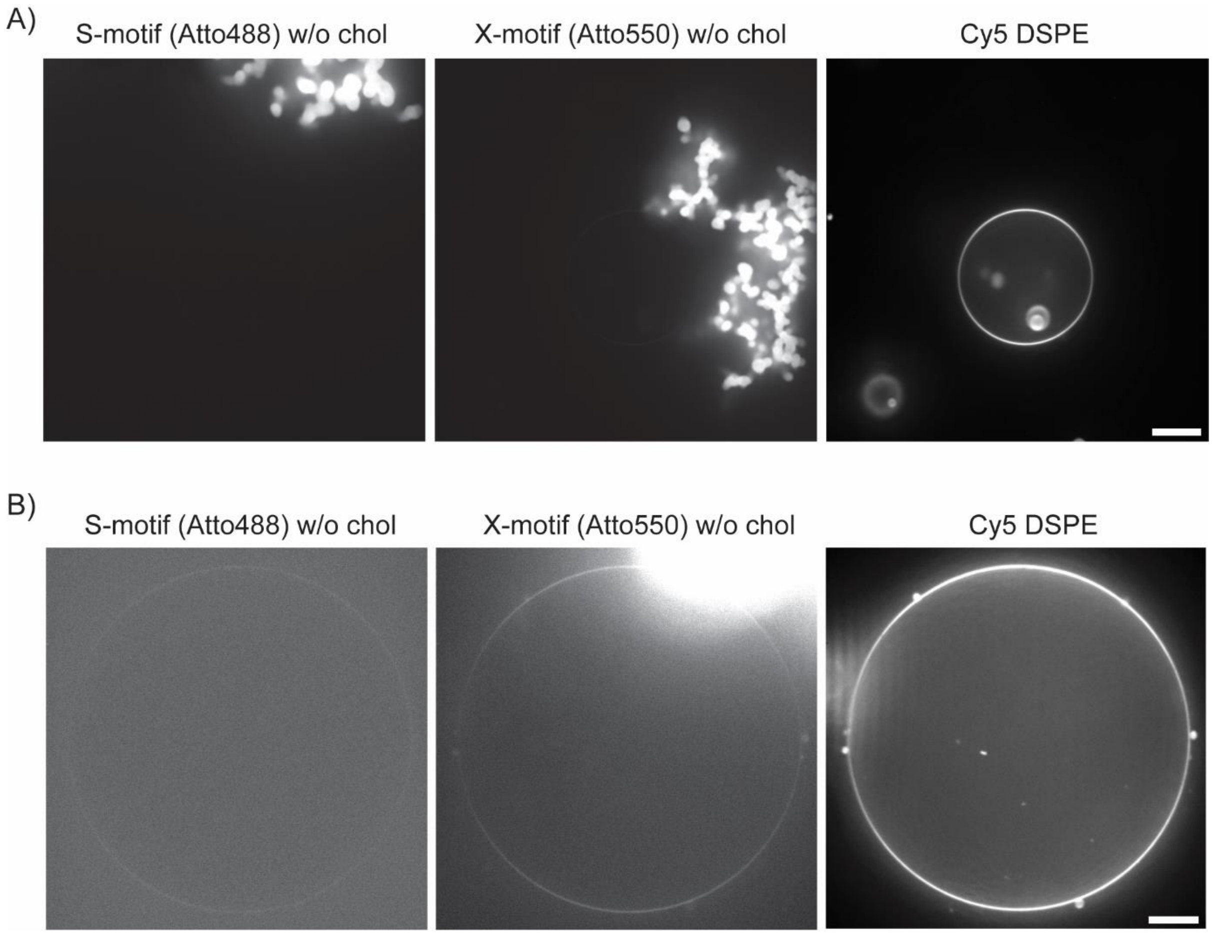

**Figure S4**: Cryo-electron microscopy images of large unilamellar vesicles with or without 75 nM S-motifs in Tris pH 8 containing 50 mM NaCl. Scale bars: 50 nm.

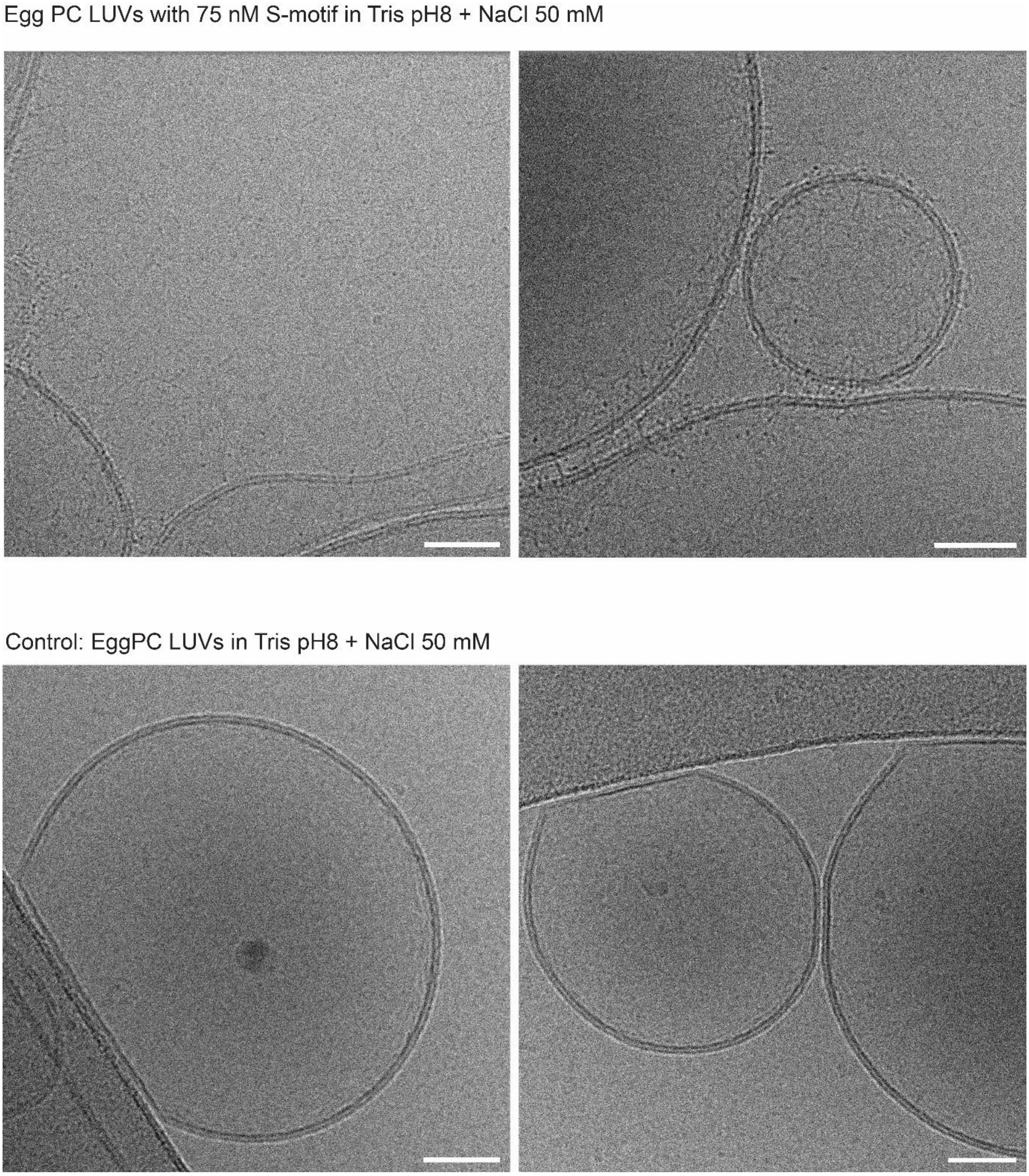

**Figure S5:** Fluorescence microscopy images of giant unilamellar vesicles (99% Egg PC with 1% Rhodamine-DHPE) with externally added S-motifs with varied DNA and NaCl concentrations. Scale bars: 10 µm.

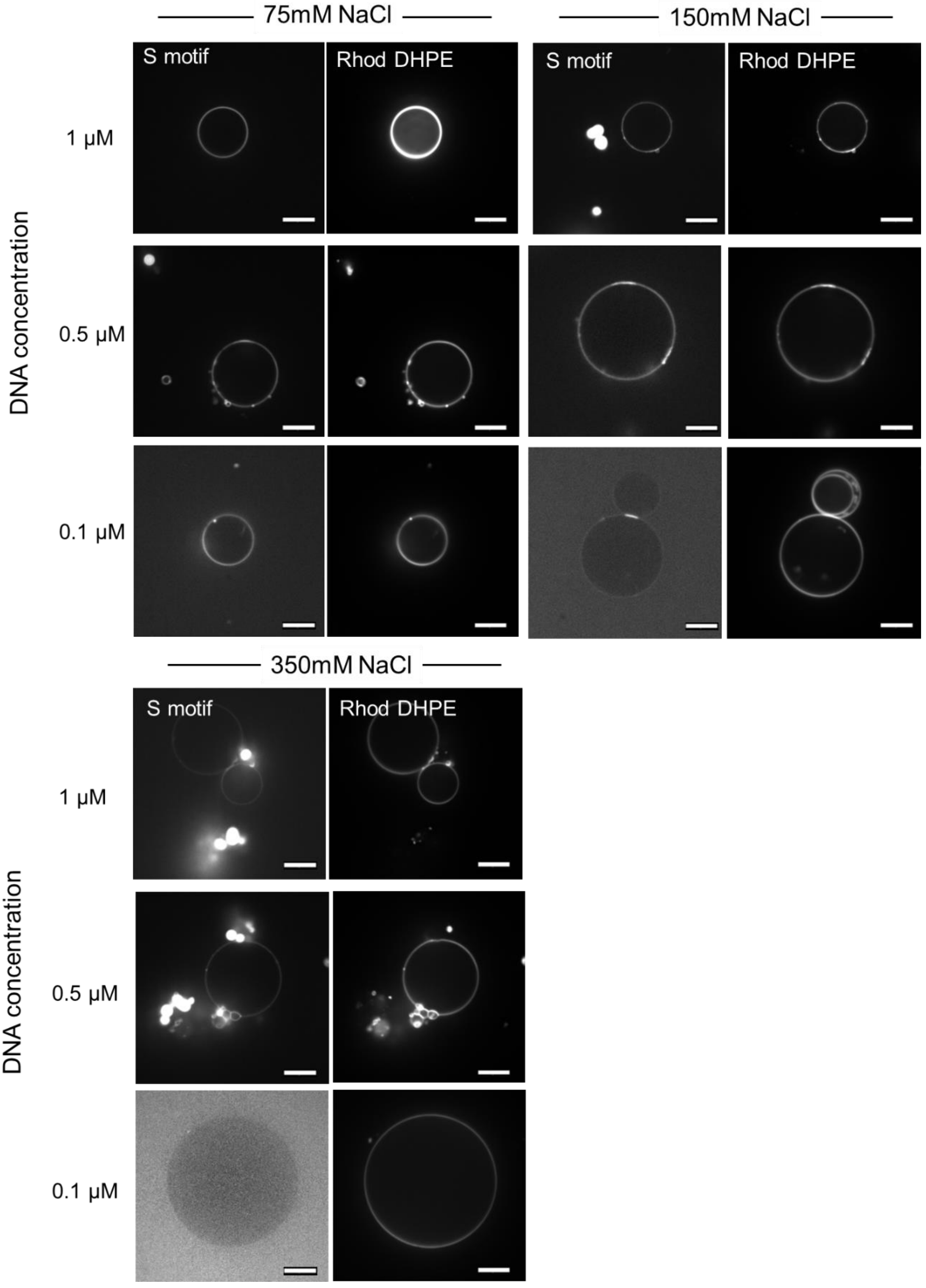

**Figure S6:** Fluorescence microscopy images of GUV (99% Egg PC, 1% Rhodamine-DHPE) co-incubated with S- and X-motifs. S-motifs have been visualized by labeling 10% of motifs with Atto488, as described in the main text. X-motifs are unlabeled fluorescently, due to the overlap of the fluorescent wavelength with Rodamine-DHPE.

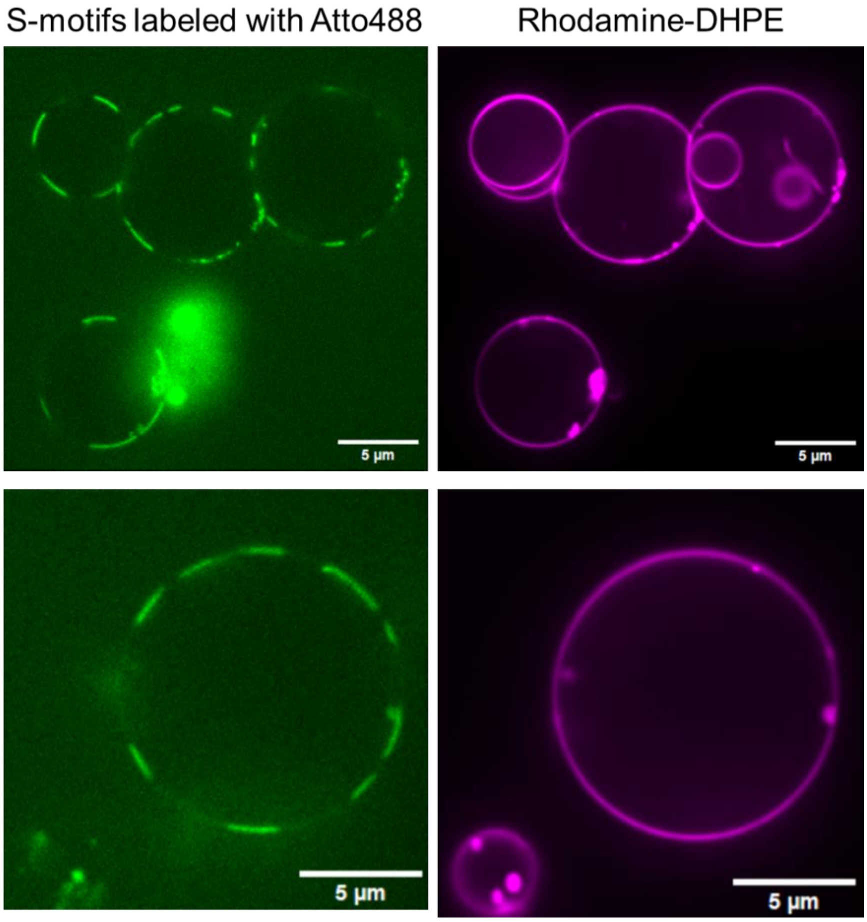

**Figure S7**: Individual fluorescence recovery after bleaching (FRAP) curves for membrane-anchored S- and X-motifs at each condition; partial bleaching of a domain and full-bleaching of an isolated domain.

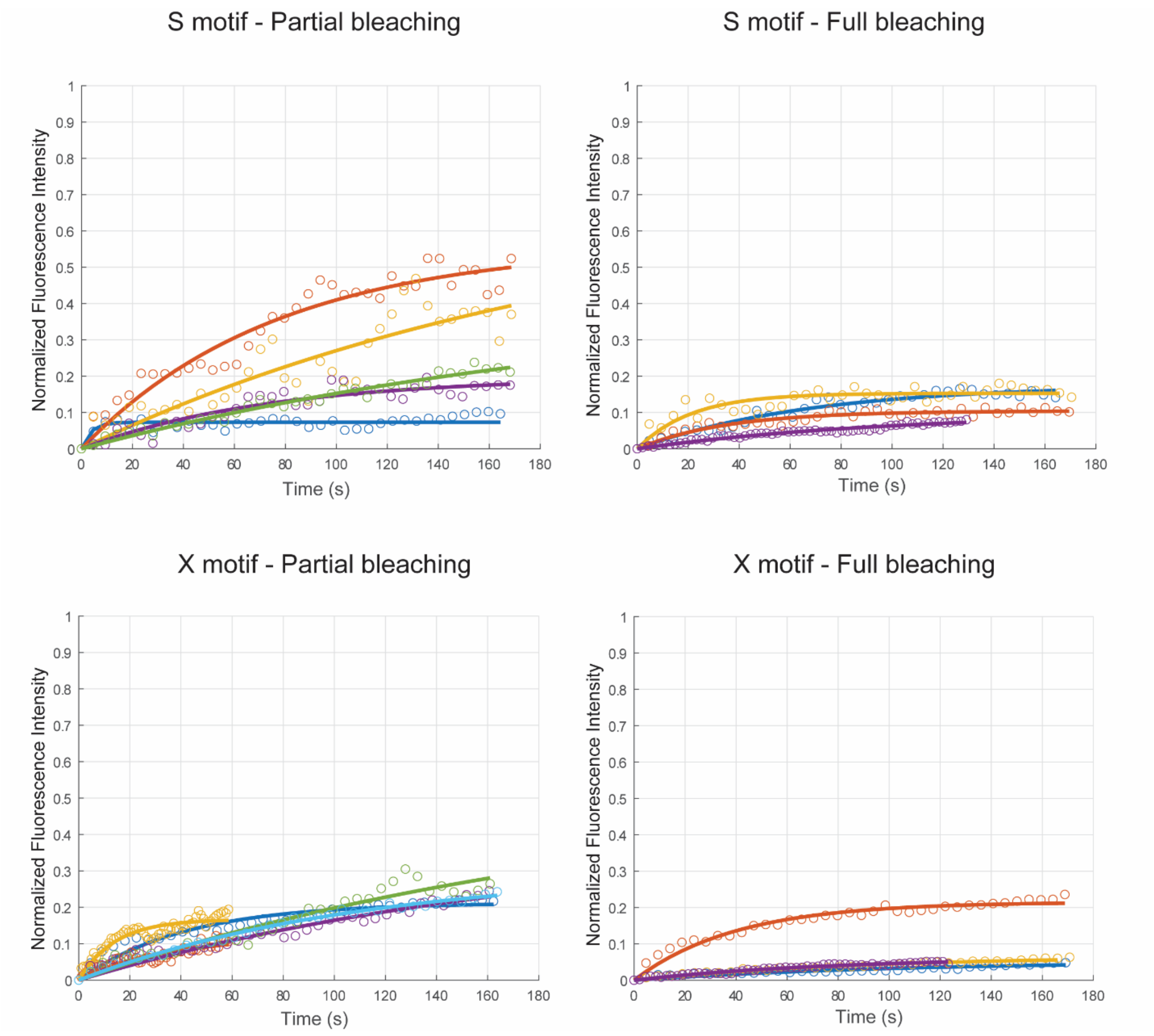

**Figure S8**: Representative images of each condition in phase diagram displayed in **Fig. 4A** in main text.

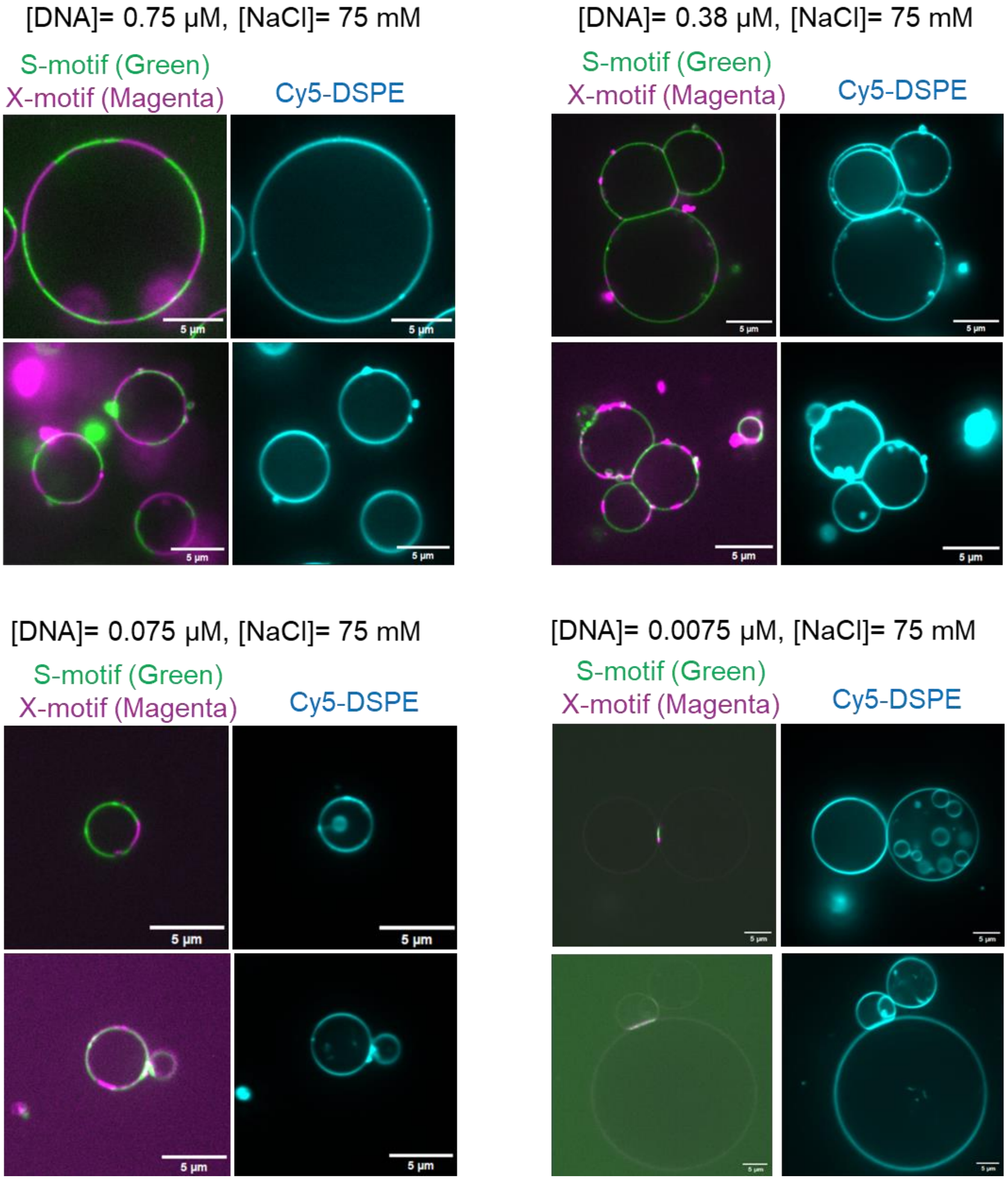

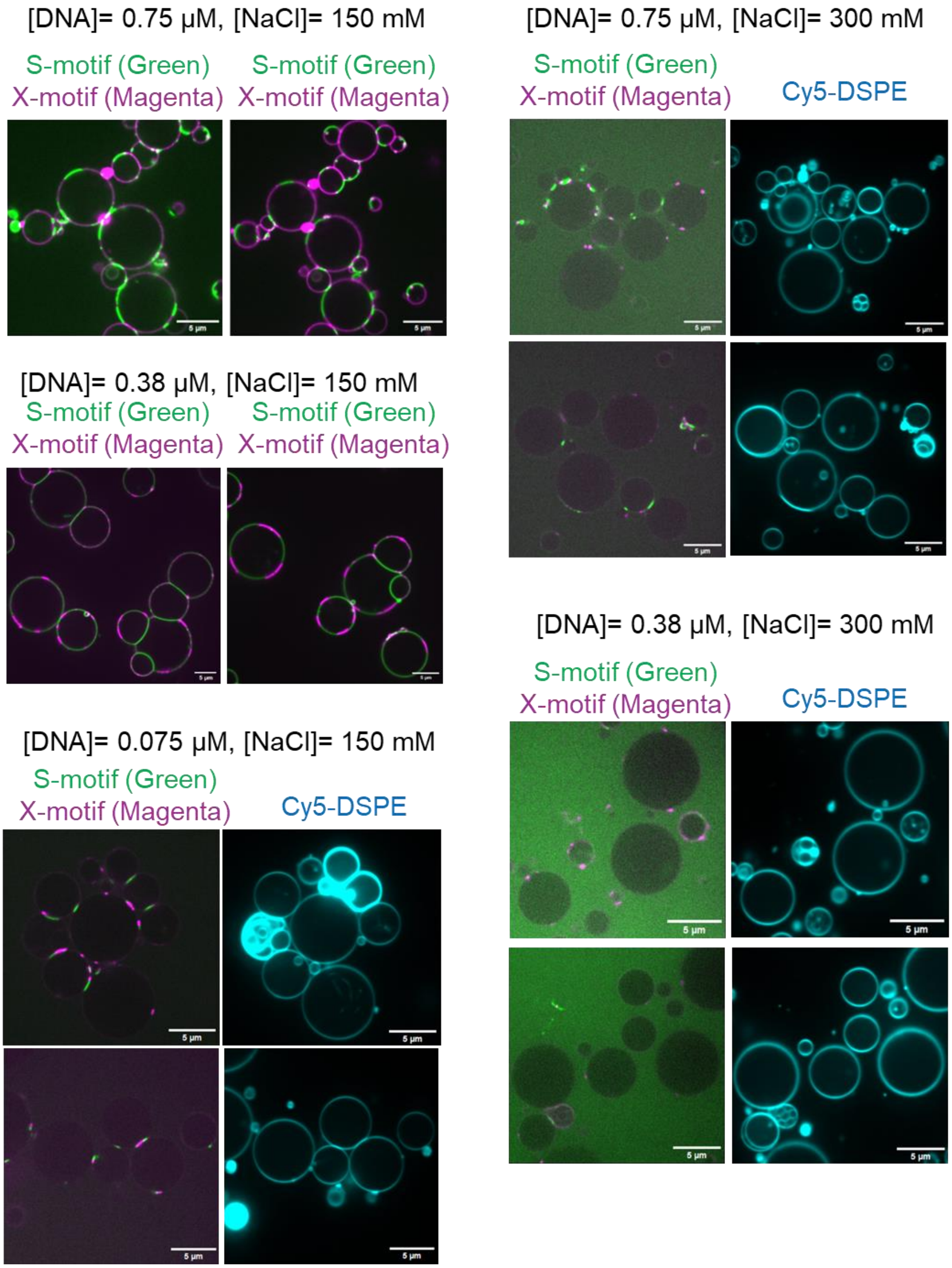
[DNA]= 0.75 µM, [NaCl]= 150 mM
S-motif (Green)
X-motif (Magenta)
S-motif (Green)
X-motif (Magenta)
[DNA]= 0.38 µM, [NaCl]= 150 mM
S-motif (Green)
X-motif (Magenta)
S-motif (Green)
X-motif (Magenta)
[DNA]= 0.075 µM, [NaCl]= 150 mM
S-motif (Green)
X-motif (Magenta)
Cy5-DSPE
[DNA]= 0.75 µM, [NaCl]= 300 mM
S-motif (Green)
X-motif (Magenta)
Cy5-DSPE
[DNA]= 0.38 µM, [NaCl]= 300 mM
S-motif (Green)
X-motif (Magenta)
Cy5-DSPE
